# The Dual-Antigen Ad5 COVID-19 Vaccine Delivered as an Intranasal Plus Subcutaneous Prime Elicits Th1 Dominant T-Cell and Humoral Responses in CD-1 Mice

**DOI:** 10.1101/2021.03.22.436476

**Authors:** Adrian Rice, Mohit Verma, Annie Shin, Lise Zakin, Peter Sieling, Shiho Tanaka, Joseph Balint, Kyle Dinkins, Helty Adisetiyo, Brett Morimoto, Wendy Higashide, Justin Taft, Rosheel Patel, Sofija Buta, Marta Martin-Fernandez, Dusan Bogunovic, Patricia Spilman, Elizabeth Gabitzsch, Jeffrey T. Safrit, Shahrooz Rabizadeh, Kayvan Niazi, Patrick Soon-Shiong

**Affiliations:** ImmunityBio, Inc., 9920 Jefferson Blvd, Culver City, CA 90232, USA; Center for Inborn Errors of Immunity, Icahn School of Medicine at Mount Sinai, 1 Gustave Lane, Levy Place, New York, NY 10029-5674, USA; Department of Pediatrics, Icahn School of Medicine at Mount Sinai, 1 Gustave Lane, Levy Place, New York, NY 10029-5674, USA; Precision Immunology Institute, Icahn School of Medicine at Mount Sinai, 1 Gustave Lane, Levy Place, New York, NY 10029-5674, USA; Mindich Child Health and Development Institute, Icahn School of Medicine at Mount Sinai, 1 Gustave Lane, Levy Place, New York, NY 10029-5674, USA; Department of Microbiology, Icahn School of Medicine at Mount Sinai, 1 Gustave Lane, Levy Place, New York, NY 10029-5674, USA

**Author notes:** These authors contributed equally (Adrian Rice, Mohit Verma).

## Abstract

In response to the need for an efficacious, thermally-stable COVID-19 vaccine that can elicit both humoral and cell-mediated T-cell responses, we have developed a dual-antigen human adenovirus serotype 5 (hAd5) COVID-19 vaccine in formulations suitable for subcutaneous (SC), intranasal (IN), or oral delivery. The vaccine expresses both the SARS-CoV-2 spike (S) and nucleocapsid (N) proteins using an hAd5 platform with E1, E2b, and E3 sequences deleted (hAd5[E1-, E2b-, E3-]) that is effective even in the presence of hAd5 immunity. In the vaccine, S is modified (S-Fusion) for enhanced cell-surface display to elicit humoral responses and N is modified with an Enhanced T-cell Stimulation Domain (N-ETSD) to direct N to the endosomal/lysosomal pathway to increase MHC I and II presentation. Initial studies using subcutaneous (SC) prime and SC boost vaccination of CD-1 mice demonstrated that the hAd5 S-Fusion + N-ETSD vaccine elicits T-helper cell 1 (Th1) dominant T-cell and humoral responses to both S and N. We then compared SC to IN prime vaccination with either an SC or IN boost post-SC prime and an IN boost after IN prime. These studies reveal that IN prime/IN boost is as effective at generating Th1 dominant humoral responses to both S and N as the other combinations, but that the SC prime with either an IN or SC boost elicits greater T cell responses. In a third study to assess the power of the two routes of delivery when used together, we used a combined SC plus IN prime with or without a boost and found the combined prime alone to be as effective as the combined prime with either an SC or IN boost in generating both humoral and T-cell responses. The findings here in CD-1 mice demonstrate that combined SC and IN prime-only delivery has the potential to provide broad immunity – including mucosal immunity – against SARS-CoV-2 and supports further testing of this delivery approach in additional animal models and clinical trials.

## INTRODUCTION

In response to the need for a COVID-19 vaccine that is safe, effective, and suitable for global distribution, we have developed the dual antigen hAd5 S-Fusion + N-ETSD vaccine including formulations for subcutaneous (SC), oral, and intranasal (IN) delivery. The vaccine comprises the SARS-CoV-2 spike (S) protein optimized for cell surface expression (S-Fusion) ^1^ to increase humoral responses and the nucleocapsid protein with an Enhanced T-cell Stimulation Domain (N-ETSD) to target N to the endosomal/lysosomal cellular compartment to enhance MHC I and II presentation.^2^

The vaccine antigens are delivered using a human adenovirus serotype 5 (hAd5) vector with deletions in the E1, E2b, and E3 gene regions (hAd5 [E1-, E2b-, E3-]; Supplementary Fig. S1A).^3^ Specifically, removal of the E2b regions confers advantageous immune properties by minimizing immune responses to Ad5 viral proteins such as viral fibers, ^4^ thereby eliciting potent immune responses to specific antigens in patients with pre-existing adenovirus (Ad) immunity. ^5,6^ Since these deletions allow the hAd5 platform to be efficacious even in the presence of existing Ad immunity, this platform enables relatively long-term antigen expression without significant induction of anti-vector immunity. It is therefore also possible to use the same vector/construct for homologous prime-boost therapeutic regimens.^7^ Importantly, this next generation Ad vector has demonstrated safety in over 125 patients with solid tumors. In these Phase I/II studies, CD4+ and CD8+ antigen-specific T cells were successfully generated to multiple somatic antigens (CEA, MUC1, brachyury) even in the presence of pre-existing Ad immunity.^5 8^

SARS-CoV-2 is an enveloped positive sense, single-strand RNA β coronavirus primarily composed of four structural proteins - spike (S), nucleocapsid (N), membrane (M), and envelope (E) – as well as the viral membrane and genomic RNA. The S glycoprotein ^9-11^ is displayed as a trimer on the viral surface, whereas N is located within the viral particle (Supplementary Fig. S1B).Spike initiates infection by the SARS-CoV-2 virus by interaction of its receptor binding domain (RBD) with the human host angiotensin-converting enzyme 2 (ACE2) expressed on the surface of cells in the respiratory system, including alveolar epithelial cells, ^12^ as well as cells in the digestive tract.

The majority of current SARS-CoV-2 vaccines under development deliver only the S antigen because antibodies raised against S RBD are expected to neutralize infection.^13–15^ Reliance on S as the sole vaccine antigen is not without risk, however, particularly in the face of the rapidly dominating variants including the B.1.351 variant expressing E484K, K417N, and N501Y mutations; ^16^ the B.1.1.7 variant (N501Y); ^17,18^ and the Cal.20.C L452R variant ^19^ all of which have altered RBD sequences that may not be as effectively recognized by antibodies generated in response to first-wave S-based vaccines.

To lessen the risk of single-antigen delivery and to broaden protective immune responses, we included the N protein in our hAd5 S-Fusion + N-ETSD vaccine (Supplementary Fig. S1C). N is a highly conserved and antigenic SARS-CoV-2-associated protein that has been studied previously as an antigen in coronavirus vaccine design for SARS-CoV.^20–23^ N associates with viral RNA and has a role in viral RNA replication, virus particle assembly, and release.^24,25^ Studies have shown that nearly all patients infected with SARS-CoV-2 have antibody responses to N.^26,27^ Furthermore, another study reported that most, if not all, COVID-19 survivors tested were shown to have N-specific CD4+ T-cell responses.^15^

The ability of N to elicit vigorous T-cell responses highlights another advantage of the addition of N. A robust T-cell response to vaccination is at least as important as the production of antibodies ^28^ and should be a critical consideration for COVID-19 vaccine efficacy. First, humoral and T-cell responses are highly correlated, with titers of neutralizing antibodies being proportional to T-cell levels, suggesting the T response is necessary for an effective humoral response.^29^ It is well established that the activation of CD4+ T helper cells enhances B-cell production of antibodies. Second, virus-specific CD4+ and CD8+ T cells are widely detected in COVID-19 patients,^30^ based on findings from patients recovered from the closely-related SARS-CoV, and there are reports that such T cells persist for at least 6–17 years, suggesting that T cells may be an important part of long-term immunity.^31–33^ These T-cell responses were predominantly to N, as described in Le Bert *et al*., who found that in all 36 convalescent COVID-19 patients in their study, the presence of CD4+ and CD8+ T cells recognizing multiple regions of the N protein could be demonstrated.^33^ They further examined blood from 23 individuals who had recovered from SARS-CoV and found that the memory T cells acquired 17 years ago also recognized multiple proteins of SARS-CoV-2. These findings emphasize the importance of designing a vaccine with the highly conserved nucleocapsid present in both SARS-CoV and SARS-CoV-2. Third, recovered patients exposed to SARS-CoV-2 have been found without seroconversion, but with evidence of T-cell responses.^34^ The T-cell based responses become even more critical given the finding in at least one study that neutralizing antibody titers decline in some COVID-19 patients after about 3 months.^35^ The importance of both S and N was highlighted by Grifoni *et al*.^15^ who identified both S and N antigens as *a priori* potential B and T-cell epitopes for the SARS-CoV virus that shows close similarity to SARS-CoV-2 that are predicted to induce both T and B cell responses.

Additional considerations for vaccine design beyond the choice of antigens, include the practicality of global distribution and the ability to generate mucosal immunity that provides the highest probability of preventing transmission. While the mRNA-based vaccines have shown excellent efficacy, ^36,37^ their requirement for extremely cold storage has presented a challenge, particularly for developing countries. Our hAd5 [E1-, E2b-, E3-] platform-based vaccine overcomes the need for super-cold storage, with the injectable and IN formulations requiring only −20°C (up to one year) or 2-8°C (up to one month) storage. The oral formulation has a further advantage of being stable at room temperature.

IN or oral vaccine delivery also offers the potential for conferring mucosal immunity. SARS-CoV-2 is a mucosal virus ^38,39^ that in most instances of infections, initiates infection by entry to the nose and mouth. Similarly, it’s most efficient route of transmission is by respiratory droplets that are then transmitted to other persons.^40^ Thus a vaccine that also elicits protective mucosal responses mediated by IgA is more likely to reduce transmission as compared to systemic, IgG-only humoral and T-cell responses. ^41^

It was our goal in the studies presented herein to confirm enhanced cell surface expression of S-Fusion as compared to S-WT (the localization of N-ETSD to endo/lysosomes is demonstrated in Sieling *et al*. 2020 ^2^) in *in vitro* studies, then assess humoral and T cell responses *in vivo* studies in CD-1 mice. In mice, first the immune responses to SC prime and SC boost vaccination were determined, then SC and IN prime delivery were compared. In a third experiment, the two routes of delivery were combined in a single boost to ascertain if together optimal immune responses could be achieved that may not necessarily be dependent upon a boost.

In all three study paradigms - SC prime with SC boost study, SC versus IN prime with boost, and combined SC plus IN prime with or without boost - immunization of CD-1 mice with the hAd5 S Fusion + N-ETSD vaccine elicited Th1 dominant, virus-neutralizing humoral responses against S. Both CD4+ and CD8+ T-cell responses to SARS-CoV-2 S and N peptide pools were also seen, with cytokine production being greater overall in response to N peptides in all studies. Potent neutralization of SARS-CoV-2 by sera from vaccinated mice in all studies was confirmed by a surrogate neutralization assay. ^42^ While all dosing paradigms produced broad immune responses, perhaps the most significant and compelling finding was that a single prime administration by combined SC and IN dosing generated immune responses that were at least as great as dosing regimens that included a boost.

## RESULTS

### S-Fusion enhances cell-surface display of conformationally-relevant spike

Before initiation of *in vivo* studies in mice, our goal of enhancing cell-surface display of S was confirmed by transfection of HEK-293T cells with hAd5 S wild type (S WT), S-WT plus N-ETSD, S-Fusion alone, and S-Fusion plus N-ETSD followed by flow cytometric analysis of anti-S receptor binding domain (S RBD) antibody binding. As shown in Supplementary Figure S2A-E, antibody binding was enhanced with S-Fusion as compared to S-WT and was found to be the highest for hAd5 S-Fusion plus N-ETSD. This enhanced cell surface expression was further confirmed by binding of recombinant ACE2, which was also the highest for hAd5 S-Fusion + N-ETSD (Supplementary Fig. S2F-H). The binding of both antibodies and ACE2 to S as expressed by the vaccine not only confirms increased surface expression, it verifies conformational relevance of the surface-displayed S protein.

### *In Vivo* Studies in Mice

#### Mice generated both anti-S and anti-N antibodies in response to SC prime, SC boost vaccination

In this initial study to test an SC hAd5 S-Fusion + N-ETSD prime followed by an SC boost, CD-1 female mice received 1 x 10^10^ viral particles (VP) of either hAd5 Null or the vaccine, both n = 5, on Days 0 and 21. Mice were euthanized and tissue collected for analysis on Day 28 (Fig. 1A).

Vaccinated mice generated anti-S antibodies that were shown to be neutralizing in surrogate and live virus assays (Fig. 1B, C, and D, respectively). Vaccinated mice also generated anti-N antibodies (Fig. 1E), and both the anti-S and anti-N antibodies were Th1 dominant (Fig. 1F).

Our neutralization data with live SARS-CoV-2 virus demonstrated the potency of the antibody response generated following vaccination with hAd5 S-Fusion + N-ETSD, with evidence of high neutralization even at a high dilution factor. In addition, a synergistic effect of pooled sera was evident, with potent neutralization even greater than control convalescent serum at ≥ 1:1,000 dilution.

**Fig. 1.**
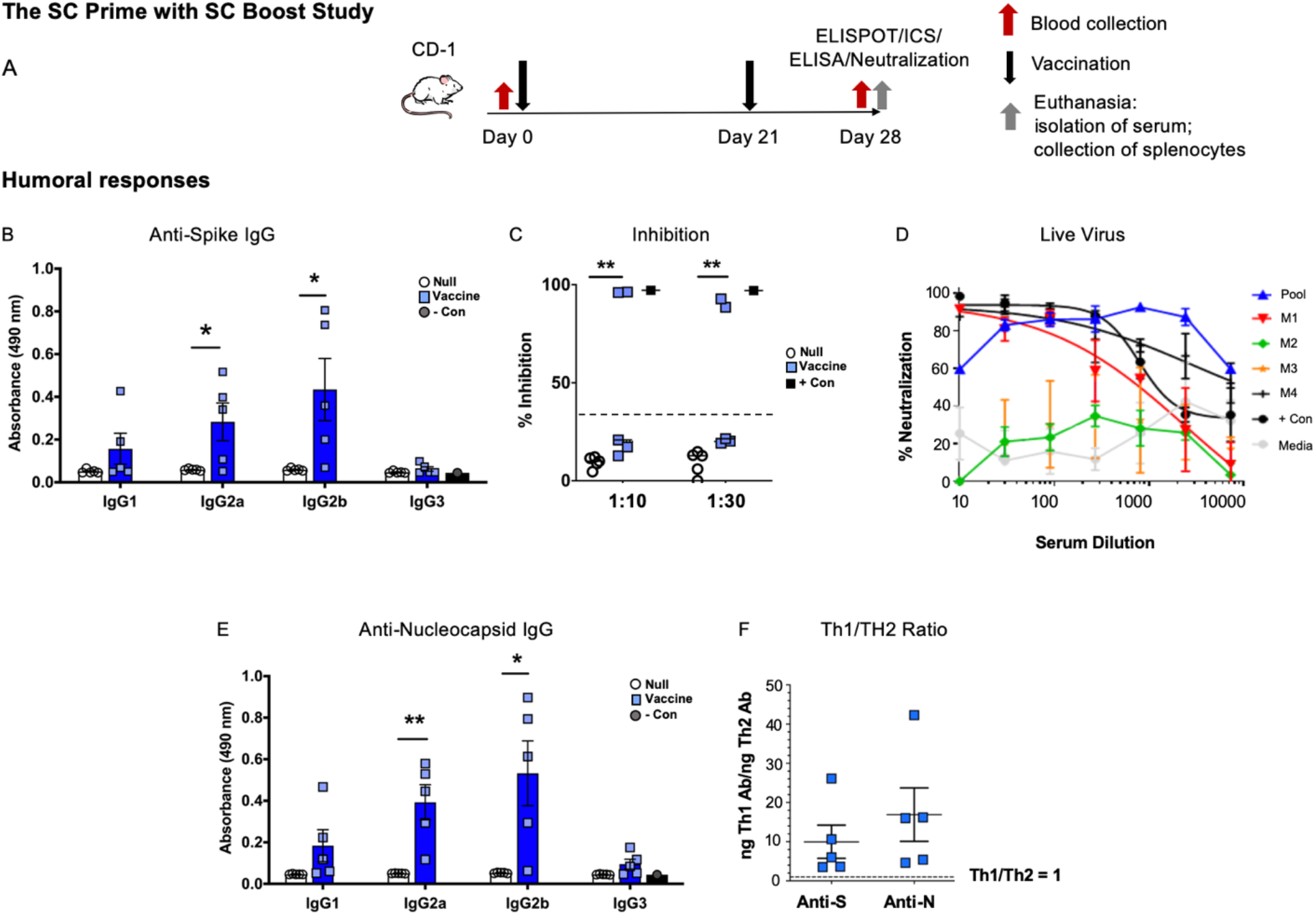
Humoral responses in the SC prime, SC boost study. (A) CD-1 mice received either hAd5 Null or the hAd5 S-Fusion + N-ETSD vaccine (both n = 5) on Day 0 and Day 21 by subcutaneous (SC) injection and were euthanized for tissue collection on Day 28. (B) Anti-spike (S) antibody levels in sera by isotype are shown (dilution 1:30); (C) results of the surrogate spike receptor binding domain (S RBD): angiotensin-converting enzyme 2 (ACE2) binding assay where inhibition of 35% or greater is associated with neutralization; and (D) percent neutralization in the live virus assay are shown for the 4 vaccinated mice with assessible sera as well as pooled sera from these mice. (E) Anti-nucleocapsid (N) antibody levels (dilution 1:90) by isotype and (F) the T helper cell 1 (Th1)/Th2 ratios for both anti-S and anti-N antibodies with a ratio greater than 1 representing Th1 dominance. Statistics performed using an unpaired two-tailed Student’s t-test where *p < 0.05 and **p ≤ 0.01.

### N peptides elicited strong CD4+ T-cell, but S peptides elicited strong CD8+ T-cell responses in SC prime, SC boost vaccinated mice

Intracellular cytokine staining (ICS) revealed that the N peptide pool stimulated interferon-γ (IFN-γ) and tumor necrosis factor alpha (TNF-α) production in selected CD4+ T-lymphocytes from vaccinated but not hAd5 Null mice (Fig. 2A and B). Conversely, the S1 peptide pool (containing S RBD) elicited higher IFN-γ/TNF-α production in CD8+ T-lymphocytes than the N peptide pool (Fig. 2C and D). ELISpot showed N peptides stimulated higher IFN-γ secretion than the S1 peptide pool, but cytokine secretion was greater with both stimuli in T-cells from vaccinated mice as compared to Null (Fig. 2E). IL-4 secretion was very low, therefore the T-cell responses, like humoral responses, were Th1 dominant with the IFN-γ/IL-4 ratio being >1 in 4 of 5 vaccinated mice (Fig. 2F and G).

**Fig. 2.**
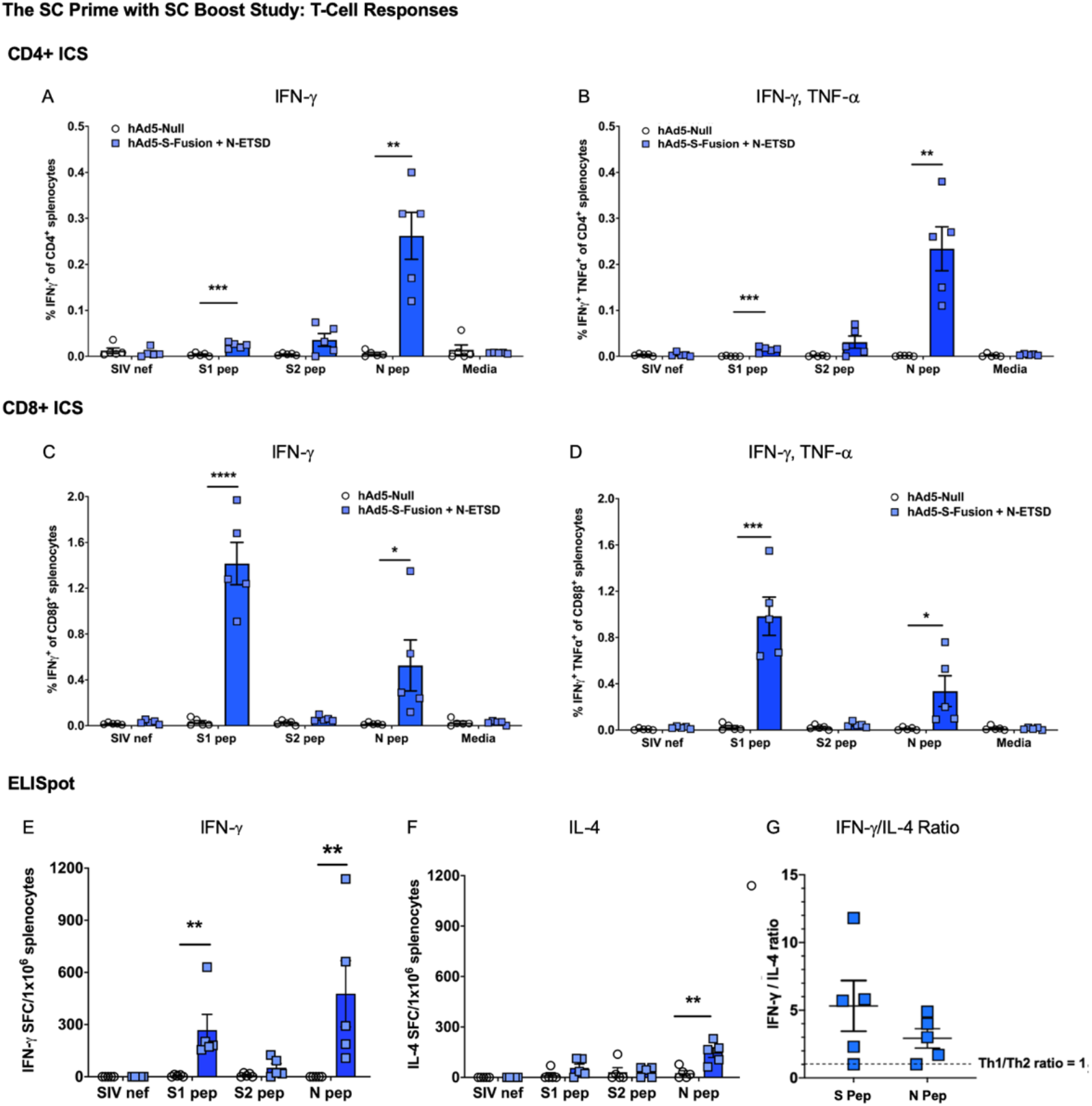
T-cell responses in SC prime, SC boost hAd5 S-Fusion + N-ETSD vaccinated mice. Production of interferon-γ (IFN-γ) for (A) CD4+ and (C) CD8β+ T lymphocytes; and IFN-γ/tumor necrosis factor α (TNF-α) by (B) CD4+ and (D) CD8β+ T lymphocytes by intracellular cytokine staining (ICS) is shown. ELISpot detection of (E) IFN-γ and (F) interleukin-4 (IL-4) by T-lymphocytes is shown. In both ICS and ELISpot, cytokine production is stimulated by exposure to S1, S2 or N peptide pools; media only and SIV nef are negative controls. (G) The IFN-γ/IL-4 ratio > 1 reflects Th1 dominance. Statistical analyses performed using an unpaired, two-tailed Student’s t-test where *p < 0.05, **p ≤ 0.01, and ***p ≤ 0.001.

### IN prime with an IN boost vaccination with hAd5 S-Fusion + N-ETSD elicited humoral responses that were as good or better than SC prime with either SC or IN boost

To both enhance and broaden immune responses by generation of mucosal immunity, we next performed a study wherein the prime delivery by either SC and intranasal (IN) routes would be compared when followed by either an SC or IN boost. The design of the SC versus IN prime with SC or IN boost study is shown in Figure 3A. There were 4 groups of CD-1 mice: untreated, SC prime followed by SC boost (SC > SC), IN prime followed by IN boost (IN > IN), and SC prime followed by IN boost (SC > IN). SC doses were administered at 1 x 10^10^ VP and IN doses were administered at 1 x 10^9^ VP. The untreated group was n = 4, SC > SC and SC > IN were n = 8 and IN > IN n =7. Mice received the priming doses on Day 0 and boosting doses on Day 21. All mice were euthanized on Day 28 and tissue including blood for serum, spleens for T cells, and lung tissue collected for analyses.

Mice in all vaccinated groups produced anti-S IgG and overall, levels were the highest in sera from IN > IN group mice (Fig. 3B). Sera were highly neutralizing as reflected by high inhibition in the surrogate virus neutralization assay (Fig. 3C). Anti-N IgG was also detected in sera from all vaccinated mice, with the levels being very similar between vaccinated groups (Fig. 3D).

Similar to the findings for sera, anti-S IgG was detected in lung homogenate of all vaccinated mice and was higher overall for the IN > IN group (Fig. 3E). Lung homogenate from all IN > IN group mice showed high inhibition in the surrogate neutralization assay, whereas homogenate from 4 mice in the SC > SC boost group did not surpass the 35% level of inhibition that is associated with viral neutralization (Fig. 3F). In lung homogenate, anti-N IgG showed a trend to higher in the SC > IN group (Fig. 3G). Not unexpectedly, both anti-S and anti-N IgA levels in lung homogenate were highest in the IN > IN boost group (Fig. 3H). Furthermore, the anti-S and anti-N responses in both sera and lung were highly Th1 dominant for all vaccinated groups (Fig. 3I).

**Fig. 3.**
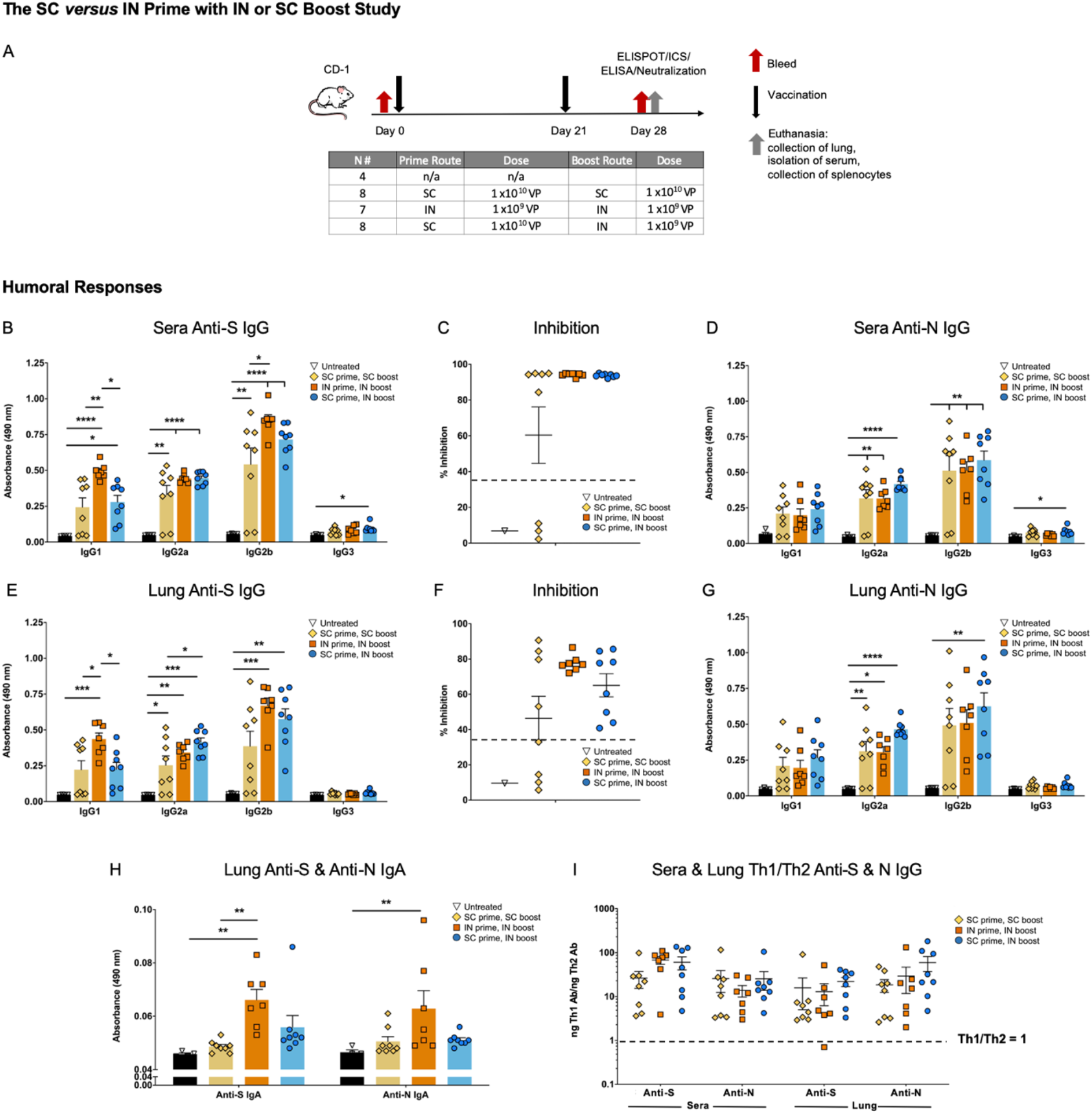
Humoral responses in the SC versus IN prime with SC or IN boost study. (A) CD-1 mice were untreated (n = 4) or received an SC prime, SC boost (n = 8); IN prime, IN boost (n = 7); or SC prime, IN boost (n = 8). (B) Sera levels of anti-S antibodies (dilution 1:30) by subtype are shown and (C) percent inhibition in the surrogate neutralization assay of ACE2:S RBD binding wherein inhibition of 35% or greater is associated with neutralization of viral infection. (D) Levels of anti-N IgG in sera (dilution 1:270). (E) Anti-S IgG by subtype (dilution 1:30) and (F) neutralization by lung homogenate. (G) Lung anti-N IgG levels (dilution 1:30). (H) Both anti-S and anti-N IgA in lung homogenate. (I) The Th1/Th2 ratios for sera and lung anti-S and anti-N antibodies where values greater than 1 represent Th1 dominance. Statistical analyses performed using One-way ANOVA with Tukey’s post-hoc analysis comparing each group to every other group where *p < 0.05; **p ≤ 0.01; ***p ≤ 0.001; and ****p ≤ 0.0001.

### Both CD4+ and CD8+ T-cell responses were greater to N than S, and higher with SC delivery

Intracellular cytokine staining (ICS) of IFN-g (Fig. 4A, D); IFN-γ, TNF-α (Fig. 4B, E), and IFN-γ, TNF-α, interleukin-2 (IL-2) (Fig. 4C, F) showed the highest mean values for the SC > SC boost and SC > IN vaccinated groups with responses to the N peptide pool trending higher for both CD4+ and CD8+ T cells. This was somewhat in contrast with the findings of the first SC > SC boost study (study 1 above) where CD8+ T-cell responses were greater to the S1 peptide pool (Fig. 2C), however, variation is expected in outbred CD-1 mice and robust CD8+ responses to both S and N were detected in SC > SC mice from each study. While the differences were not statistically significant due to variation among individual mice, overall the IN > IN boost group had a reduced population of CD8+ cells capable of accumulating IFN-γ (Fig. 4D) and IFN-γ + TNF-α (Fig. 4E) in response to S and N peptide stimulation.

ELISpot findings were similar, with higher responses seen for the SC > SC and SC > IN groups when compared to the IN > IN group, and the highest responses were found to be specific to the N peptide pool (Fig. 4G). Interleukin-4 (IL-4) secretion in ELISpot was very low for all groups (Fig. 4H), therefore the IFN-γ/IL-4 ratios were above 1 for almost all vaccinated mice in response to both S and N peptide pools (Fig. 4I).

The T-cell responses in this study suggested an important contribution of SC delivery to T cell responses.

**Fig. 4.**
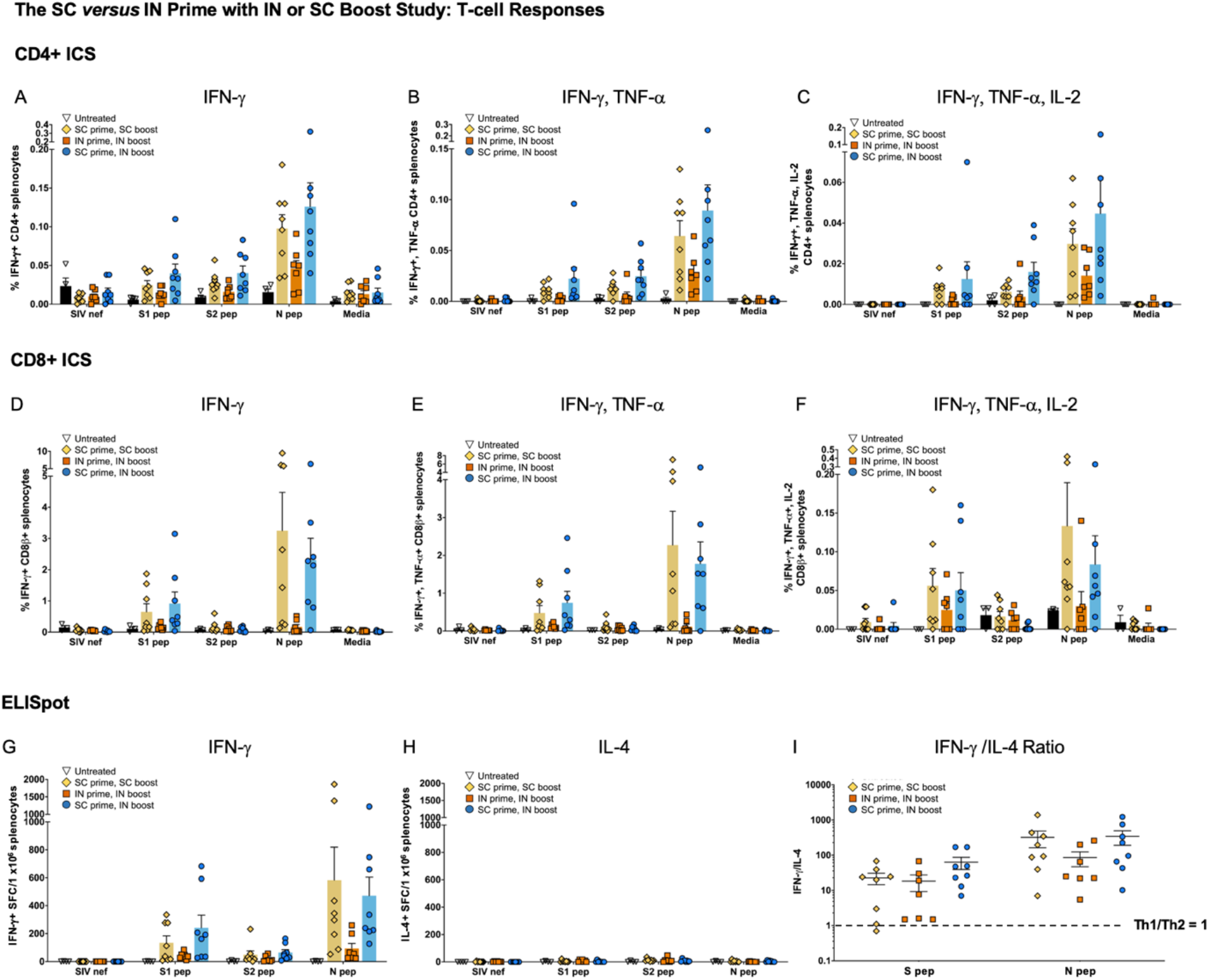
Both CD4+ and CD8+ T cells respond the nucleocapsid peptides in SC versus IN prime study with SC or IN boost. Cytokine production in response to S1, S2 and N peptide pools as detected by Intracellular cytokine staining (ICS) of (A) Interferon-γ (IFN-γ); (B) IFN-γ and tumor necrosis factor α (TNF-α); and (C) IFN-γ, TNF-α, and interleukin-2 (IL-2) production by CD4+ is shown. ICS (D-F) IFN-γ; IFN-γ, TNF-α; and IFN-γ, TNF-α, and IL-2 for CD8+ T cells is shown. Some outliers by the Grubb’s test were removed. ELISpot for (G) IFN-γ and (H) interleukin-4 (IL-4) secretion in response to the peptide pools is shown. SIV nef is a negative control. (I) The IFN-γ/IL-4 ratio showing T helper cell 1 dominance. Statistical analyses performed using One-way ANOVA with Dunnett’s post-hoc comparison of each treatment group to untreated for each peptide pool was performed but did not reveal statistically significant differences due to individual variation amongst mice.

### Prime-only delivery by combined SC and IN dosing elicits humoral responses that are as good or better than those with a boost

To leverage both the humoral responses effectively elicited by IN delivery with the T-cell responses that were greater with SC delivery, we then tested prime delivery by a combination of the SC and IN routes, with either IN or SC boosts. This study design is shown in Figure 5A. There were 5 groups of CD-1 mice: untreated, an SC prime at 1 x 10^10^ viral particles (VP) without boost (SC > no boost), a combined 9 x 10^9^ VP SC plus 1 x 10^9^ VP IN prime (SC/IN) without boost (SC/IN > no boost), a combined SC/IN prime with 1 x 10^9^ VP IN boost (SC/IN > IN), and a combined SC/IN prime with a 1 x 10^10^ VP SC boost (SC/IN > SC). The untreated group was n = 4 and all vaccinated groups were n = 7. Mice received the prime on Day 0 and in appropriate groups, the boost on Day 21. All mice were euthanized on Day 35 and tissue including blood for serum, spleens for T cells, and lung tissue collected for analyses. Note this euthanasia day is one week later than the two studies described above (Figs. 1A and 3A), which was a change meant to better characterize humoral responses at a time point at which we expected cell-mediated responses to remain high based on our prior work with this vaccine platform.

The combined SC/IN > no boost regimen was just as effective in eliciting neutralizing anti-S IgG and anti-N IgG antibody production in both sera (Fig. 5B-D) and lung (Fig. 5E-G) as either the SC/IN > IN or SC/IN > SC regimens. SC > no boost gave significantly lower humoral responses (Fig. 5B-G). All humoral responses were Th1 dominant (Fig. 5H).

**Fig. 5.**
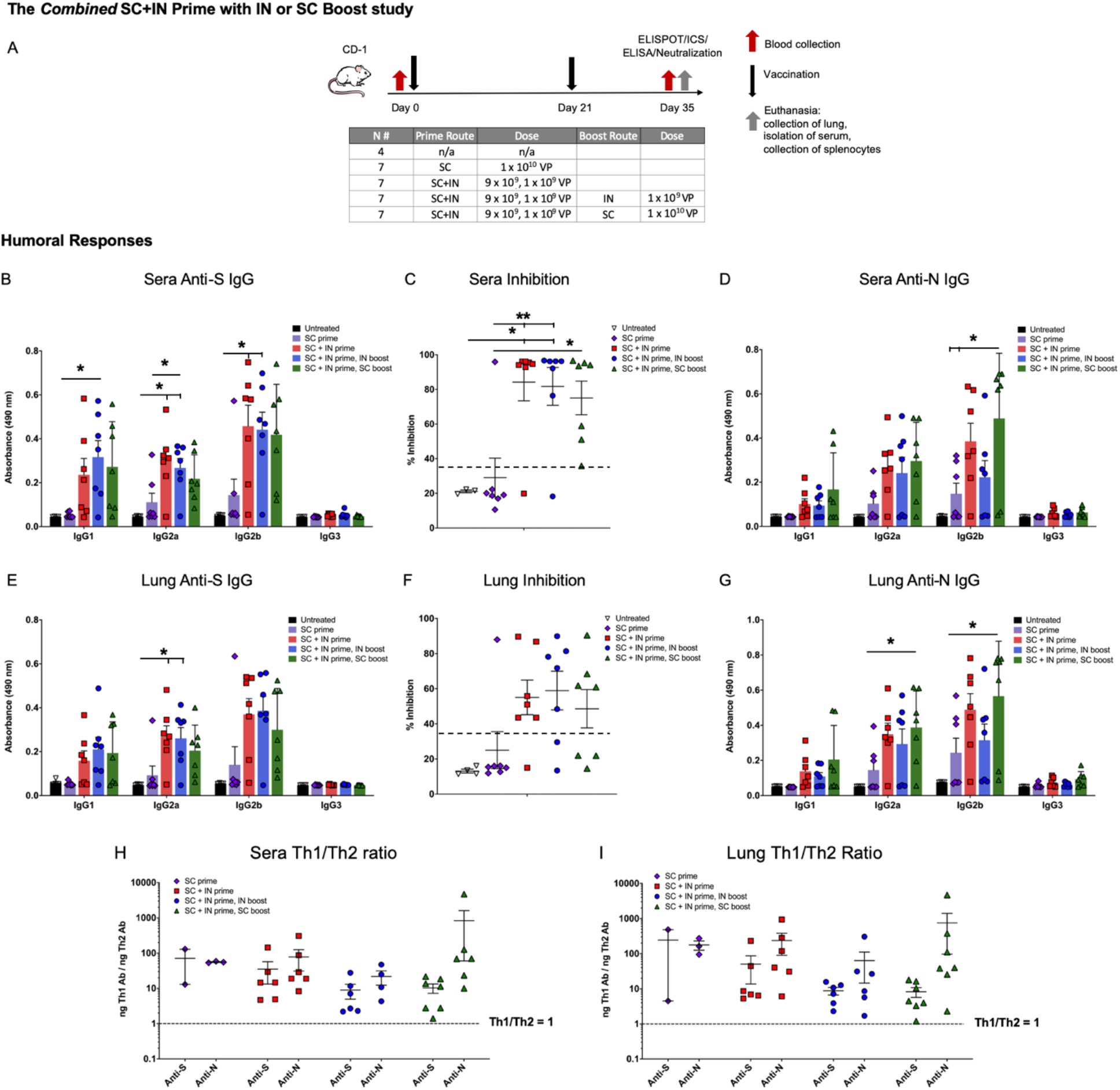
Subcutaneous (SC) plus intranasal (IN) prime without boost elicits Th1 dominant neutralizing anti-S and anti-N antibodies. (A) The study design is shown with groups for SC prime only, SC + IN prime only, and SC + IN prime with either an SC or IN boost, all n = 7. There was an untreated control group (n = 4). Prime dosing was on Day 0, boosts on Day 21, and euthanasia on Day 35. Shown are (B) serum anti-spike (S) antibodies by subtype (dilution 1:30); (C) % inhibition in the surrogate neutralization assay with sera (>35% is correlated with neutralization of virus); and (D) serum anti-nucleocapsid (N) antibodies (dilution 1:270). The same readouts (E) anti-S antibodies; (F) neutralization; and (G) anti-N antibodies are shown for lung homogenate (dilution 1:30 for anti-S and −N). The Th1/Th2 ratios for anti-S and anti-N antibodies are shown for (H) sera and (I) lung. Statistical analyses performed using One-way ANOVA with Tukey’s post-hoc analysis comparing treatment groups within each antibody subtype or groups in the neutralization assay where *p < 0.05 and ** p ≤ 0.01.

### SC plus IN prime alone without a boost elicits CD4+ T cell responses to N and CD8+ T-cell responses to S

Similar to the findings in the first study, ICS shows the N peptide pool stimulated cytokine production by CD4+ T lymphocytes from all vaccinated mice (Fig. 6A-C), but CD8+ T cells from vaccinated mice responded to S peptide pool 1 which contains the S RBD (Fig. 6D-F). The differences between vaccinated groups were not significant due to variability amongst mice, with SC/IN > no boost vaccinated mice having T-cell responses that were similar to those seen with mice that did receive boosts.

In ELISpot, the highest IFN-γ secretion in response to peptide pools differed by both peptide pool and vaccination regimen. As compared to the negative control (SIV nef), T cell IFN-γ secretion was significantly greater for the combined SC/IN > SC group in response to the S1 peptide pool; greater for the SC/IN > IN group to the S2 peptide pool; and greater for the SC/IN > no boost group to the N peptide pool (Fig. 6G).

IL-4 secretion was again very low (Fig. 6H), therefore the IFN-γ/IL-4 ratio was above 1 for all vaccinated mice with only one exception, reflecting Th1 dominance of T-cell responses (Fig. 6I).

**Fig. 6.**
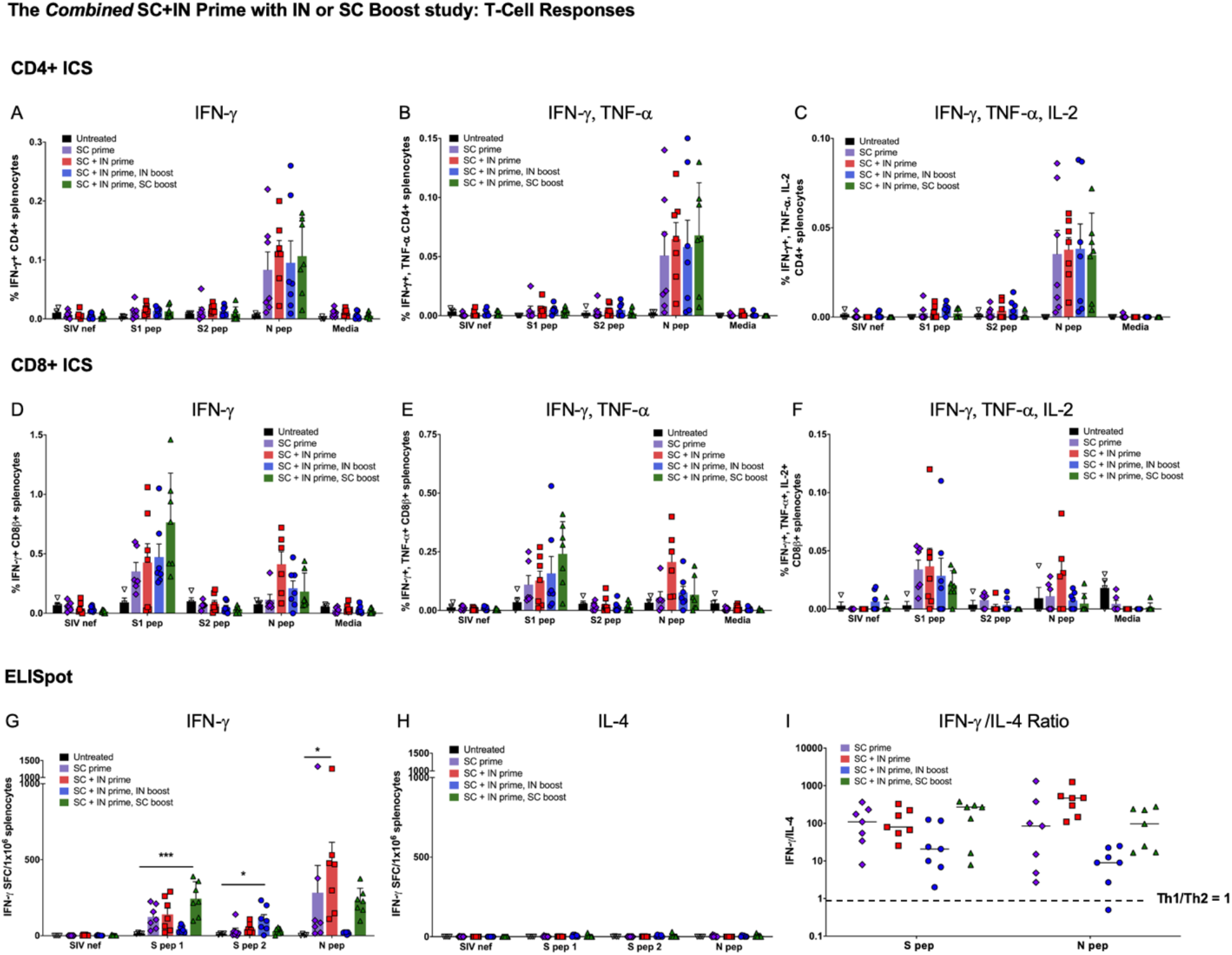
CD4+ T-cells respond to nucleocapsid (N) and CD8+ T cells to the spike in the combined SC plus IN prime study with SC or IN boost. Interferon-γ (IFN-γ); IFN-γ and tumor necrosis factor a (TNF-α); and (C) IFN-γ, TNF-α, and interleukin-2 (IL-2) production by CD4+ (A-C) and CD8+ (D-F) T-cells, respectively, in response to spike 1 (S1, containing the receptor binding domain), S2, and N peptide pools as detected by intracellular cytokine staining (ICS). Some outliers by the Grubb’s test were removed. ELISpot for (G) IFN-γ and (H) interleukin-4 (IL-4) secretion in response to the peptide pools. SIV nef and media are negative controls. (I) The IFN-γ/IL-4 ratio showing T helper cell 1 dominance. Statistical analyses performed using One-way ANOVA with Dunnett’s post-hoc comparison of each treatment group to untreated for each peptide pool where *p < 0.05, **p ≤ 0.01, and ***p ≤ 0.001.

## DISCUSSION

Our hAd5 S-Fusion + N-ETSD vaccine was designed to overcome the risks of an S-only vaccine and elicit both T-cell immunity and neutralizing antibodies, leveraging the vital role T cells play in generating long-lasting antibody responses and in directly killing infected cells. The CD4+ and CD8+ T cells responses induced by this vaccine are multifunctional, and induction of such multifunctional T cells by vaccines is correlated with better protection against infection.^43^ We posit that enhanced CD4+ T-cell responses and Th1 predominance resulting from expression of an S antigen optimized for surface display and an N antigen optimized for endosomal/lysosomal subcellular compartment localization and thus MHC I and II presentation, led to increased dendritic cell presentation, cross-presentation, B cell activation, and ultimately high neutralization capability. Furthermore, the potent neutralization capability at high dilution seen for sera from hAd5 S-Fusion + N-ETSD vaccinated mice, combined with Th1 dominance of antibodies generated in response to both S and N antigens, supports the objective of this vaccine design.

It is well established that the contemporaneous MHC I and MHC II presentation of an antigen by the antigen presenting cell activates CD4+ and CD8+ T cells simultaneously and is optimal for the generation of memory B and T cells. A key finding of our construct is that N-ETSD, which we show is directed to the endosomal/lysosomal compartment, elicits a CD4+ response, a necessity for induction of memory T cells and helper cells for B cell antibody production. Others have also reported on the importance of lysosomal localization for eliciting the strongest T-cell IFN-γ and CTL responses, compared to natural N.^44,45^

The T-cell responses to the S and N antigens expressed by hAd5 S-Fusion + N-ETSD were polycytokine, including IFN-γ, TNF-α, and IL-2 consistent with successful antimicrobial immunity in bacterial and viral infections.^46–50^ Post-vaccination polycytokine T-cell responses have been shown to correlate with vaccine efficacy, including those with a viral vector.^43^ Highly relevant here, polycytokine T-cell responses to SARS-CoV-2 N protein are consistent with recovered COVID-19 patients,^20^ suggesting that the bivalent hAd5 S-Fusion + N-ETSD vaccine will provide vaccine subjects with greater protection against SARS-CoV-2.

The key finding here that prime-only vaccination delivered by combination SC and IN dosing results in broad humoral and T-cell responses, with the potential for enhanced mucosal immunity, supports the ongoing clinical testing of the hAd5 S-Fusion + N-ETSD. The vaccine has currently completed Phase 1 testing as an SC prime and SC boost, and oral boost formulations that have shown efficacy in the ability to elicit both humoral and T-cell responses that conferred complete protection against high-titer SARS-CoV-2 challenge in our pre-clinical studies in non-human primates, ^1^ will soon also be tested in the clinic. To our knowledge, our vaccine is currently the only one available in SC, thermally-stable oral,^51^ and IN formulations that offer the expanded possibilities for efficient, feasible delivery across the globe, particularly in developing nations.

## ACKNOWLEDGEMENTS

We thank Phil Yang of ImmunityBio for his ongoing coordination of project reports for this study.

## SUPPLEMENTARY INFORMATION

### METHODS

#### The hAd5 [E1-, E2b-, E3-] platform and constructs

For studies here, the next generation hAd5 [E1-, E2b-, E3-] vector was used (Fig. S1A) to create viral vaccine candidate constructs. This hAd5 [E1-, E2b-, E3-] vector is primarily distinguished from other first-generation [E1-, E3-] recombinant Ad5 platforms ^52,53^ by having additional deletions in the early gene 2b (E2b) region that remove the expression of the viral DNA polymerase (pol) and in pre terminal protein (pTP) genes, and its propagation in the E.C7 human cell line.^3,4,54,55^

The hAd5 S-Fusion + N-ETSD vaccine we utilized the hAd5 [E1-, E2b-, E3-] comprises a wildtype spike (S) sequence [accession number YP009724390] modified with a proprietary linker peptide sequence as well as a wildtype nucleocapsid (N) sequence [accession number YP009724397] with a an Enhanced T-cell Stimulation Domain (ETSD) signal sequence to direct translated N to the endosomal/lysosomal pathway. ^2^ The SARS-CoV-2 S protein is found on the viral surface ^11^ and the N protein is found in the interior of the virus ^23,56^ (Fig. S1B).

The powerful cytomegalovirus (CMV) promoter ^57^ drives expression in the hAd5 construct (Fig. S1C).

**Fig. S1.**
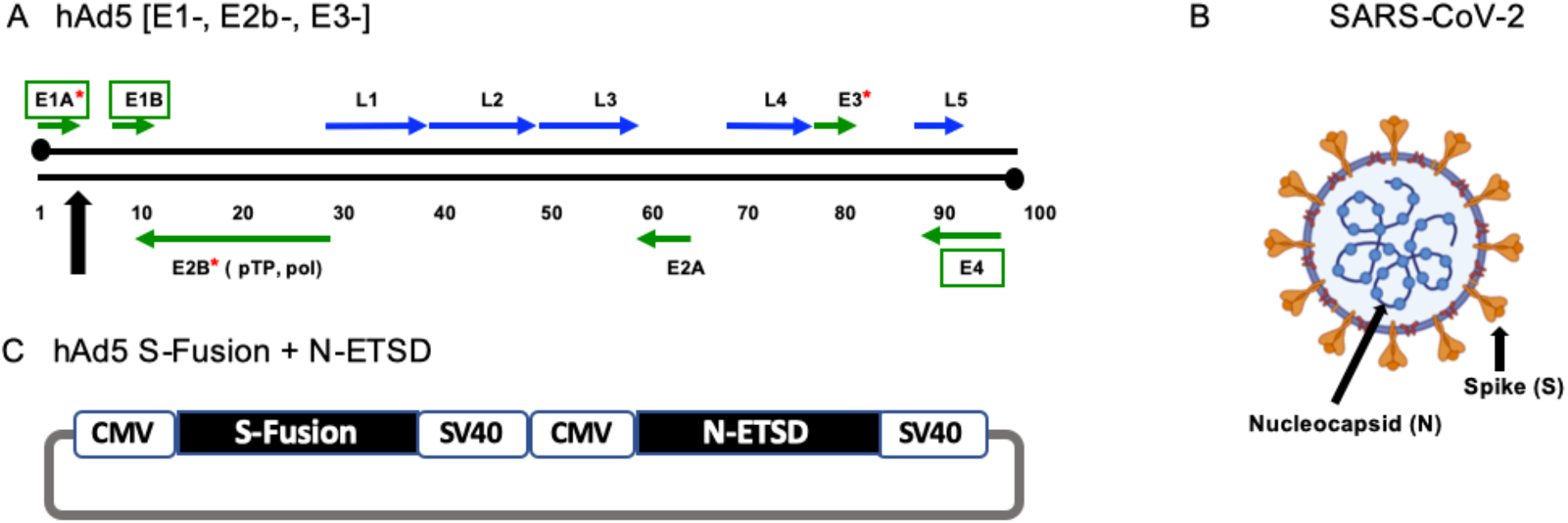
The SARS-CoV-2 virus, the hAd5 [E1-, E2b-, E3-] vector and the dual antigen hAd5 S-Fusion + N-ETSD vaccine. (A) The second-generation human adenovirus serotype 5 (hAd5) vector used has the E1, E2b, and E3 regions deleted. Sequences for the vaccine antigen cargo are inserted at the black arrow. (B) The spike (S) glycoprotein is displayed as a trimer on the surface of SARS-CoV-2 and the nucleocapsid (N) protein is found in the virus interior, associated with the viral RNA. (C) The vaccine antigens are under control of the cytomegalovirus (CMV) promoter and sequences end with SV40 poly-A.

#### Transfection of HEK 293T cells with hAd5 constructs and flow cytometric analysis of RBD surface expression

To determine surface expression of the RBD epitope by vaccine candidate constructs, we transfected HEK 293T cells with hAd5 construct DNA and quantified surface RBD by flow cytometric detection using anti-RBD antibodies. The constructs tested were: S-WT, S-WT + N-ETSD, S-Fusion, S-Fusion + N-ETSD, and N-ETSD. HEK 293T cells (2.5 x 10^5^ cells/well in 24 well plates) were grown in DMEM (Gibco Cat# 11995-065) with 10% FBS and 1X PSA (100 units/mL penicillin, 100 μg/mL streptomycin, 0.25 ug/mL Amphotericin B) at 37°C. Cells were transfected with 0.5 μg of hAd5 plasmid DNA using a JetPrime transfection reagent (Polyplus Catalog # 89129-924) according to the manufacturer’s instructions. Cells were harvested 1, 2, 3, and 7 days post transfection by gently pipetting cells into medium and labeled with an anti-RBD monoclonal antibody (clone D003 Sino Biological Catalog # 40150-D003) and F(ab’)2-Goat anti-Human IgG-Fc secondary antibody conjugated with R-phycoerythrin (ThermoFisher Catalog # H10104). Labeled cells were acquired using a Thermo-Fisher Attune NxT flow cytometer and analyzed using Flowjo Software.

#### ACE2-IgG1Fc binding to hAd5 transfected HEK 293T cells

HEK 293T cells were cultured at 37°C under conditions described above for transfection with hAd5 S-WT, S-Fusion, and S-Fusion + N-ETSD and were incubated for 2 days and harvested for ACE2-Fc binding analysis. Recombinant ACE2-IgG1Fc protein was produced using Maxcyte transfection in CHO-S cells that were cultured for 14 days. ACE2-IgG1Fc was then purified using a MabSelect SuRe affinity column on AKTA Explorer. Purified ACE2-IgG1Fc was dialyzed into 10 mM HEPES, pH7.4, 150 mM NaCl and concentrated to 2.6 mg/mL. For binding studies, the ACE2-IgG1Fc was used at a concentration of 1 μg/mL for binding. Cells were incubated with ACE2-Fc for 20 minutes and, after a washing step, were then labeled with a PE conjugated F(ab’)2-goat anti-human IgG Fc secondary antibody at a 1:100 dilution, incubated for 20 minutes, washed and acquired on flow cytometer. Histograms are based on normalized mode (NM) of cell count – count of cells positive for signal in PE channel.

#### Murine immunization and blood/tissue collection

CD-1 female mice (Charles River Laboratories) 6-8 weeks of age were used for immunological studies performed at the vivarium facilities of Omeros Inc. (Seattle, WA). Mice were administered subcutaneous (SC) injections at the indicated doses in 50 μL ARM buffer (20 mM Tris pH 8.0, 25 mM NaCl, with 2.5% glycerol) or intranasal (IN) injections at the indicated doses in 10 μL ARM buffer (5 μL per nostril) while under isoflurane anesthesia. On the final day of each study, blood was collected via the submandibular vein from isoflurane-anesthetized mice for isolation of sera using a microtainer tube and then mice were euthanized for collection of spleen and lungs.

Spleens were removed from each mouse and placed in 5 mL of sterile media (RPMI/HEPES/Pen/Strep/10% FBS). Splenocytes were isolated within 2 hours of collection and used fresh or frozen for later analysis.

Lungs were removed from each mouse, dissected in half and then immediately snap frozen on dry ice. Lung homogenates were generated by thawing one frozen lung half and homogenizing in 150 μL sterile PBS using a Fisher Scientific pestle drill. Homogenates were centrifuged at 13,000 rpm for 3 minutes and supernatants were utilized in ELISA and cPass surrogate neutralization assays.

#### Intracellular cytokine stimulation (ICS)

ICS assays were performed using 10^6^ live splenocytes per well in 96-well U-bottom plates. Splenocytes in RPMI media supplemented with 10% FBS were stimulated by the addition of pools of overlapping peptide for S or N antigens at 2 μg/mL/peptide for 6 h at 37°C in 5% CO_2_, with protein transport inhibitor, GolgiStop (BD) added two hours after initiation of incubation. The S peptide pool (JPT: Cat #PM-WCPV-S-1) is a total of 315 spike peptides split into two pools comprised of 158 and 157 peptides each. The N peptide pool (JPT; Cat # PM-WCPV-NCAP-1) was also used to stimulate cells. A SIV-Nef peptide pool (BEI Resources) was used as an off-target negative control. Stimulated splenocytes were then stained for a fixable cell viability stain followed by the lymphocyte surface markers CD8β and CD4, fixed with CytoFix (BD), permeabilized, and stained for intracellular accumulation of IFN-γ, TNF-α and IL-2 (in studies 2 and 3). Fluorescent-conjugated antibodies against mouse CD8β antibody (clone H35-17.2, ThermoFisher), CD4 (clone RM4-5, BD), IFN-γ (clone XMG1.2, BD), TNF-α (clone MP6-XT22, BD) and IL-2 (clone JES6-5H4; BD), and staining was performed in the presence of unlabeled anti-CD16/CD32 antibody (clone 2.4G2; BD). Flow cytometry was performed using a Beckman-Coulter Cytoflex S flow cytometer and analyzed using Flowjo Software.

#### ELISpot assay

ELISpot assays were used to detect cytokines secreted by splenocytes from inoculated mice. Fresh splenocytes were used on the same day, as were cryopreserved splenocytes containing lymphocytes. The cells (2-4 x 10^5^ cells per well of a 96-well plate) were added to the ELISpot plate containing an immobilized primary antibodies to either IFN-γ or IL-4 (BD), and were exposed to various stimuli (e.g. control peptides, target peptide pools/proteins) comprising 2 μg/mL peptide pools or 10 μg/mL protein for 36-40 hours. After aspiration and washing to remove cells and media, extracellular cytokine was detected by a secondary antibody to cytokine conjugated to biotin (BD). A streptavidin/horseradish peroxidase conjugate was used detect the biotin-conjugated secondary antibody. The number of spots per well, or per 2-4 x 10^5^ cells, was counted using an ELISpot plate reader. A Th1/Th2 ratio was calculated by dividing the IFN-γ spot forming cells (SFC) per million splenocytes with the IL-4 SFC per million splenocytes for each animal.

#### ELISA for detection of antibodies

For IgG antibody detection in sera and lung homogenate from inoculated mice, ELISAs specific for spike and nucleocapsid antibodies, as well as for IgG subtype (IgG1, IgG2a, IgG2b, and IgG3) antibodies were used. In addition, for IgA antibody detection in lung homogenate from inoculated mice, ELISAs specific for spike and nucleocapsid antibodies, as well as for IgA was used. A microtiter plate was coated overnight with 100 ng of either purified recombinant SARS-CoV-2 S-FTD (full-length S with fibritin trimerization domain, constructed and purified in-house by ImmunityBio), SARS-CoV-2 S RBD (Sino Biological, Beijing, China; Cat # 401591-V08B1-100) or purified recombinant SARS-CoV-2 nucleocapsid (N) protein (Sino Biological, Beijing, China; Cat # 40588-V08B) in 100 μL of coating buffer (0.05 M Carbonate Buffer, pH 9.6). The wells were washed three times with 250 μL PBS containing 1% Tween 20 (PBST) to remove unbound protein and the plate was blocked for 60 minutes at room temperature with 250 μL PBST. After blocking, the wells were washed with PBST, 100 μL of either diluted serum or diluted lung homogenate samples were added to wells, and samples incubated for 60 minutes at room temperature. After incubation, the wells were washed with PBST and 100 μL of a 1/5000 dilution of anti-mouse IgG HRP (GE Health Care; Cat # NA9310V), or anti-mouse IgG1 HRP (Sigma; Cat # SAB3701171), or anti-mouse IgG_2a_ HRP (Sigma; Cat # SAB3701178), or anti-mouse IgG2b HRP (Sigma; catalog# SAB3701185), anti-mouse IgG3 HRP conjugated antibody (Sigma; Cat # SAB3701192), or anti-mouse IgA HRP conjugated antibody (Sigma; Cat # A4789) was added to wells. For positive controls, a 100 μL of a 1/5000 dilution of rabbit anti-N IgG Ab or 100 μL of a 1/25 dilution of mouse anti-S serum (from mice immunized with purified S antigen in adjuvant) were added to appropriate wells. After incubation at room temperature for 1 hour, the wells were washed with PBS-T and incubated with 200 μL o-phenylenediamine-dihydrochloride (OPD substrate (Thermo Scientific Cat # A34006) until appropriate color development. The color reaction was stopped with addition of 50 μL 10% phosphoric acid solution (Fisher Cat # A260-500) in water and the absorbance at 490 nm was determined using a microplate reader (SoftMax^®^Pro, Molecular Devices).

#### Calculation of relative ng amounts of antibodies and the Th1/Th2 ratio

A standard curve of IgG was generated and absorbance values were converted into mass equivalents for both anti-S and anti-N antibodies. Using these values, we were able to calculate that hAd5 S-Fusion + N-ETSD vaccination generated a geometric mean value for S- and N-specific IgG per milliliter of serum. These values were also used to generate a Th1/Th2 ratio for the humoral responses by dividing the sum total of Th1 skewed antigen-specific IgG isotypes (IgG2a, IgG2b and IgG3) with the total Th2 skewed IgG3, for each mouse. Some responses, particularly for anti-N responses in IgG2a and IgG2b (both Th1 skewed isotypes), were above the limit of quantification with OD values higher than those observed in the standard curve. These data points were reduced to values within the standard curve, and thus will reflect a lower Th1/Th2 skewing than would otherwise be reported.

#### cPass^™^ neutralizing antibody detection

The GenScript cPass™ (https://www.genscript.com/cpass-sars-cov-2-neutralization-antibody-detection-Kit.html) for detection of neutralizing antibodies was used according to the manufacturer’s instructions.^58^ The kit detects circulating neutralizing antibodies against SARS-CoV-2 that block the interaction between the S RBD with the ACE2 cell surface receptor. It is suitable for all antibody isotypes and appropriate for use with in animal models without modification.

#### Vero E6 cell neutralization assay

All aspects of the assay utilizing virus were performed in a BSL3 containment facility according to the ISMMS Conventional Biocontainment Facility SOPs for SARS-CoV-2 cell culture studies. Vero e6 kidney epithelial cells from *Cercopithecus aethiops* (ATCC CRL-1586) were plated at 20,000 cells/well in a 96-well format and 24 hours later, cells were incubated with antibodies or heat inactivated sera previously serially diluted in 3-fold steps in DMEM containing 2% FBS, 1% NEAAs, and 1% Pen-Strep; the diluted samples were mixed 1:1 with SARS-CoV-2 in DMEM containing 2% FBS, 1% NEAAs, and 1% Pen-Strep at 10,000 TCID50/mL for 1 hr. at 37°C, 5% CO_2_. This incubation did not include cells to allow for neutralizing activity to occur prior to infection. The samples for testing included sera from the four mice that showed anti-S IgG2a and IgG2b antibody responses in Fig. 1B, pooled sera from those four mice, sera from a COVID-19 convalescent patient, and media only. For detection of neutralization, 120 μL of the virus/sample mixture was transferred to the Vero E6 cells and incubated for 48 hours before fixation with 4% PFA. Each well received 60 μL of virus or an infectious dose of 600 TCID50. Control wells including 6 wells on each plate for no virus and virus-only controls were used. The percent neutralization was calculated as 100-((sample of interest-[average of “no virus”])/[average of “virus only”])*100) with a stain for CoV-2 Np imaged on a Celigo Imaging Cytometer (Nexcelom Bioscience).

## SUPPLEMENTARY RESULTS

### Enhanced HEK 293T cell-surface expression of RBD following transfection with Ad5 S-Fusion + N-ETSD

As shown in Figure S2, anti-RBD-specific antibodies did not detect RBD on the surface of HEK 293T cells transfected with hAd5 S-WT (Fig. 2A) or hAd5 S-WT + N-ETSD (Fig. 2B) constructs, while hAd5 S-Fusion alone was higher (Fig. 2C). Notably, the highest cell-surface expression of RBD was detected after transfection with dual antigen hAd5 S-Fusion + N-ETSD (Fig. 2D). Similar results were seen for recombinant ACE2-Fc binding to S-WT, S-Fusion and S-Fusion + N-ETSD, with ACE2 showing higher binding to S-Fusion than S-WT and the dual antigen construct showing the highest binding (Fig. S2F-H). These findings support our proposition that an hAd5 S-Fusion + N-ETSD construct, containing a high number and variety of antigens provided by both full-length, optimized S with proper folding and N leads to enhanced expression and cell surface display of RBD in a vaccine construct.

**Fig. S2.**
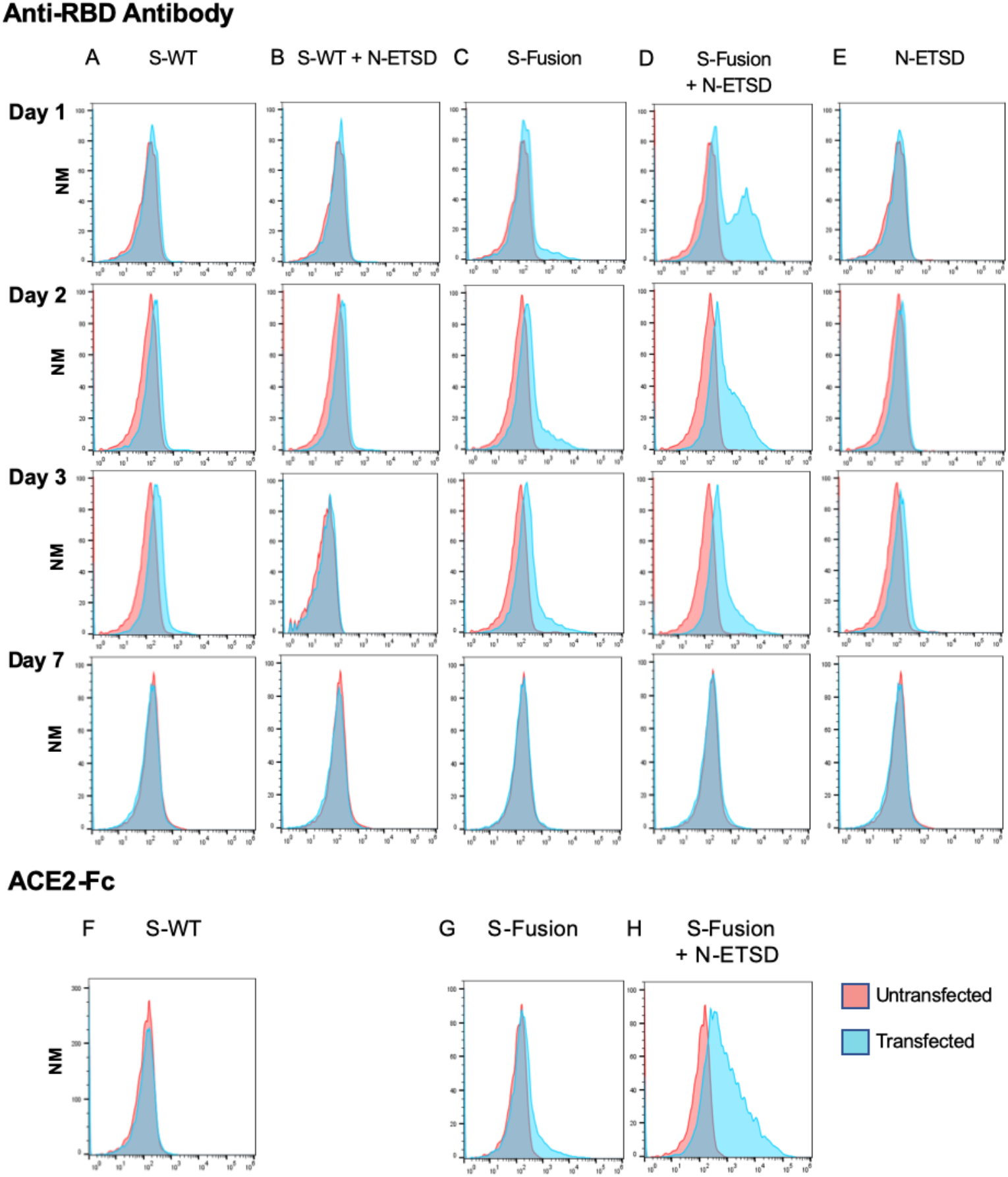
Transfection of HEK293T cells with hAd5 S-Fusion + ETSD results in enhanced surface expression of the spike receptor binding domain (RBD). Flow cytometric analysis of an anti-RBD antibody with construct-transfected cells reveals no detectable surface expression of RBD in either (A) S-WT or (B) S-WT + N-ETSD transfected cells. Expression was higher in (C) S-Fusion transfected cells as compared to S-WT. Cell surface expression of the RBD was high in (D) S-Fusion + N-ETSD transfected cells, particularly at day 1 and 2. (E) No expression was detected the N-ETSD negative control. Recombinant ACE2-Fc binding to (F) S-WT, (G) and S-Fusion, and (H) S-Fusion + N-ETSD is shown. Y-axis scale is normalized to mode (NM).

